# Deficient FGF signaling in the developing peripheral retina disrupts ciliary margin development and causes aniridia

**DOI:** 10.1101/443416

**Authors:** Revathi Balasubramanian, Chenqi Tao, Karina Polanco, Jian Zhong, Fen Wang, Liang Ma, Xin Zhang

## Abstract

The mammalian ciliary margin is a part of the developing peripheral neural retina that differentiates into the ciliary body and the iris. Canonical WNT signaling plays a critical role in the specification of the ciliary margin at the peripheral retina in the presence of strong FGF signaling in the central retina. The mechanism of how the boundary between the central retina and the ciliary margin is created has not been previously elucidated. Using genetic ablation and epistasis experiments, we show that loss of FGF signaling gradient in the peripheral retina causes expansion of WNT signaling towards the central retina thereby disrupting the neurogenic boundary and compartmentalization of the ciliary margin. Loss of WNT signaling displays a complimentary effect with expansion of FGF signaling into the ciliary marginal space. Using *in vivo* experiments, we elucidate the FGF signaling cascade involved in development of the ciliary margin. We also identify the surface ectoderm as the source of WNT ligands in eliciting WNT response at the ciliary margin. We show that an interaction between FGF and WNT signaling is required for generation of the ciliary marginal cells. Taken together, our results reveal that a gradient intersection of FGF and WNT signaling is required for specification of the ciliary margin.

## INTRODUCTION

The early development of various tissues in the body is orchestrated by individual morphogens or by the interaction of several morphogens. Typically, an interaction between two or more signaling cues is responsible for the compartmentalization within a developing structure. Development of the eye exemplifies the interactions between various signaling cues to specify the various ocular structures such as the lens, retina, retinal pigment epithelium (RPE) and ciliary margin. Eye development begins as an evagination of the developing diencephalon and initially forms the optic vesicle. The optic vesicle then reaches forward towards the surface ectoderm and invaginates, leading to the formation of the optic cup. The optic cup neuroepithelium differentiates into two distinct layers: an outer retinal pigment epithelium and an inner neural retina (Chow and Lang, 2001; Fuhrmann, 2010). The peripheral tip of the retina closest to the lens is defined as the ciliary margin. The developing ciliary margin differentiates into two important anterior segment structures – the ciliary body and the iris (Beebe, 1986; Chow and Lang, 2001; Martinez-Morales et al., 2004; Napier and Kidson, 2007). At early developmental stages, the ciliary margin at the peripheral tip of the retina is a highly proliferative zone (Fischer et al., 2013). As development proceeds, the ciliary margin is further specified and attains molecular characteristics that can be distinguished from the neural retina using specific markers (Bao and Cepko, 1997; Thut et al., 2001; Zhao et al., 2002). Prior research has shown that the ciliary margin at embryonic day 13.5 (E13.5) can be divided into proximal (closest to the neural retina), medial and distal (closest to the RPE) ciliary margin zones (Rowan et al., 2004). More recently, additional molecular markers that mark the specific compartments have been identified (Trimarchi et al., 2009). Recent studies have also shown that a population of cells in the developing ciliary margin are also capable of producing retinal neuronal cells and have a distinct molecular signature compared to retinal progenitor cells (Belanger et al., 2017; Marcucci et al., 2016). Therefore, deciphering the molecular mechanisms governing development of the ciliary margin is key to understanding normal development of the retinal neurons, ciliary body and iris.

Early work on eye development has established that FGF signaling is critical for proliferation of the optic cup progenitor cells, potentiation of neural retinal cell differentiation and inhibition of retinal pigment epithelial cell differentiation (Atkinson-Leadbeater et al., 2014; Cai et al., 2010; Guillemot and Cepko, 1992; Ou et al., 2013; Pittack et al., 1997; Sakaguchi et al., 1997). Explant studies in the chick optic vesicle showed that an exogenous addition of FGF to the developing optic vesicle causes the presumptive RPE to switch to a neural retinal fate thereby causing the formation of a double retina (Dias da Silva et al., 2007). In our previous studies using mouse models in which FGF receptors were conditionally ablated in the central neural retina, optic cups develop an RPE-like structure ectopically in the FGF signaling deficient areas (Cai et al., 2013). The activity of FGF signaling is prominent in the developing eye-cup (E8 onwards) but is later restricted to the neural retinal cells at E10.5 (Nguyen and Arnheiter, 2000). In an explant study, when FGF4 is over-expressed in the developing chick RPE, tissue abutting the ectopic source of FGF trans-differentiates into neural retina. Curiously enough, a ciliary margin forms at the junctional zone between the trans-differentiated retina and the RPE at a distance away from the FGF source (Dias da Silva et al., 2007). This indicates that FGF signaling may also be implicated in the development of the ciliary margin. A constitutive deletion of FGF9 also affects the specification of the developing ciliary margin zones (Zhao et al., 2001). Despite these findings, the role of FGF signaling in specifying the ciliary margin at the peripheral part of the neural retina has remained unexplored.

WNT signaling via the canonical β-catenin pathway is also crucial to the specification of RPE during early oculogenesis (Bharti et al., 2012; Fujimura et al., 2009; Westenskow et al., 2009). Conditional ablation of WNT/β-catenin signaling during early oculogenesis causes trans-differentiation of RPE into neural retina at E13.5. While the WNT reporter activity is high in the RPE during the early stages of optic cup development, as development proceeds, it becomes restricted to the ciliary margin (E13.5) (Ha et al., 2012; Liu et al., 2006; Trimarchi et al., 2009). Recent work has shown that the WNT/β-catenin signaling is necessary for development of the ciliary margin and an over-activation of WNT signaling in the peripheral retina leads to an ectopic expansion of the ciliary margin (Liu et al., 2007). Moreover, *in vitro* studies using mouse embryonic stem cells have shown that a dynamic regulation of WNT and FGF signaling is needed for the generation of ciliary margin like structure (Kuwahara et al., 2015). Given these findings, we seek to understand the interactions between FGF and WNT signaling during the development of the eye as it pertains to the specification of the ciliary margin.

## RESULTS

### Conditional deletion of FGF receptors in the peripheral retina causes ciliary margin defect

We conditionally deleted FGF receptor 1 (FGFR1) and FGF receptor 2 (FGFR2) using Pax6 α-Cre, also known as α-Cre, which is strongly active in the peripheral aspect of the developing retina. The loss of FGF receptors 1 and 2 results in a gradual thinning of the ciliary marginal zone starting at E11.5 (Figure 1 A, arrows). The disruption in ciliary margin morphology is especially apparent at E13.5, a developmental time point at which we have chosen to conduct our analyses going forward. The disruption in the developing ciliary margin translates to an aberrant development of the ciliary body and iris at postnatal time points (Figure 1 A). Adult mice display near complete aniridia, with the iris reduced to a short stub in the milder phenotypes (Figure 1 B), and a clump of pigmented cells with no defined architecture in more severe phenotypes (Figure 1B inset). Collectively, these data show that the loss of FGF signaling driven by α-Cre disrupts the normal development of the ciliary margin leading to aniridia.

**Figure 1:**
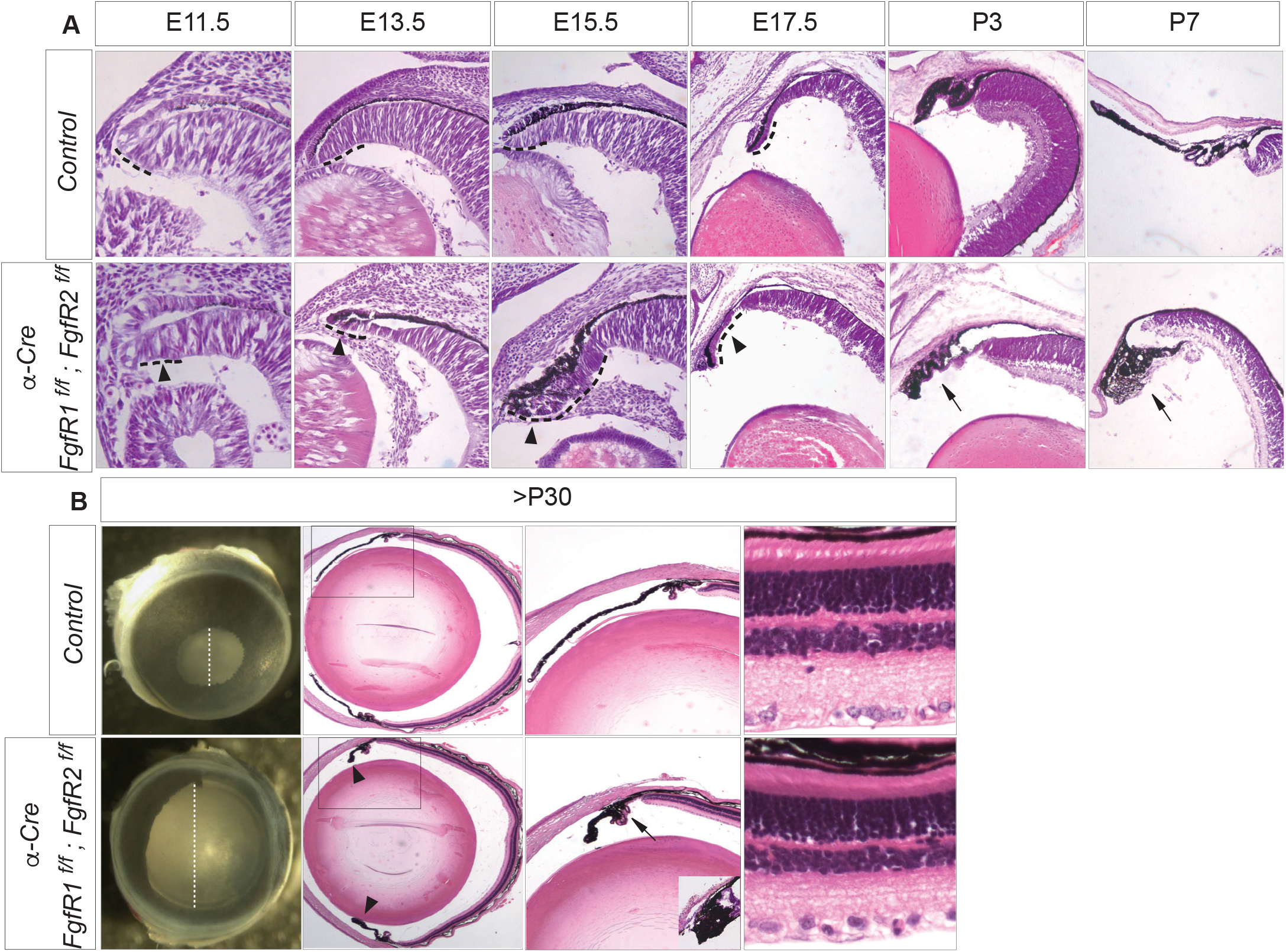
FGF signaling in the peripheral retina is required for development of the ciliary margin and iris. **A.** Loss of FGFR1 and FGFR2 in the peripheral retina leads to thinning of the ciliary margin at early developmental stages (arrowheads) and a dysmorphic ciliary body and iris at early postnatal stages (arrows). **B.** Loss of FGF signaling in the ciliary margin during development leads to iris hypoplasia. Dotted lines indicate the aberrant size of the pupil in the absence of iris.

### Loss of FGF signaling in the peripheral retina affects the compartmentalization of the ciliary margin and disrupts the neurogenic boundary

To further investigate the impact of loss of FGF signaling in the peripheral retina, we employed several retinal and ciliary margin markers to characterize the developmental disruption in compartmentalization of the ciliary margin (Figure 2).

**Figure 2:**
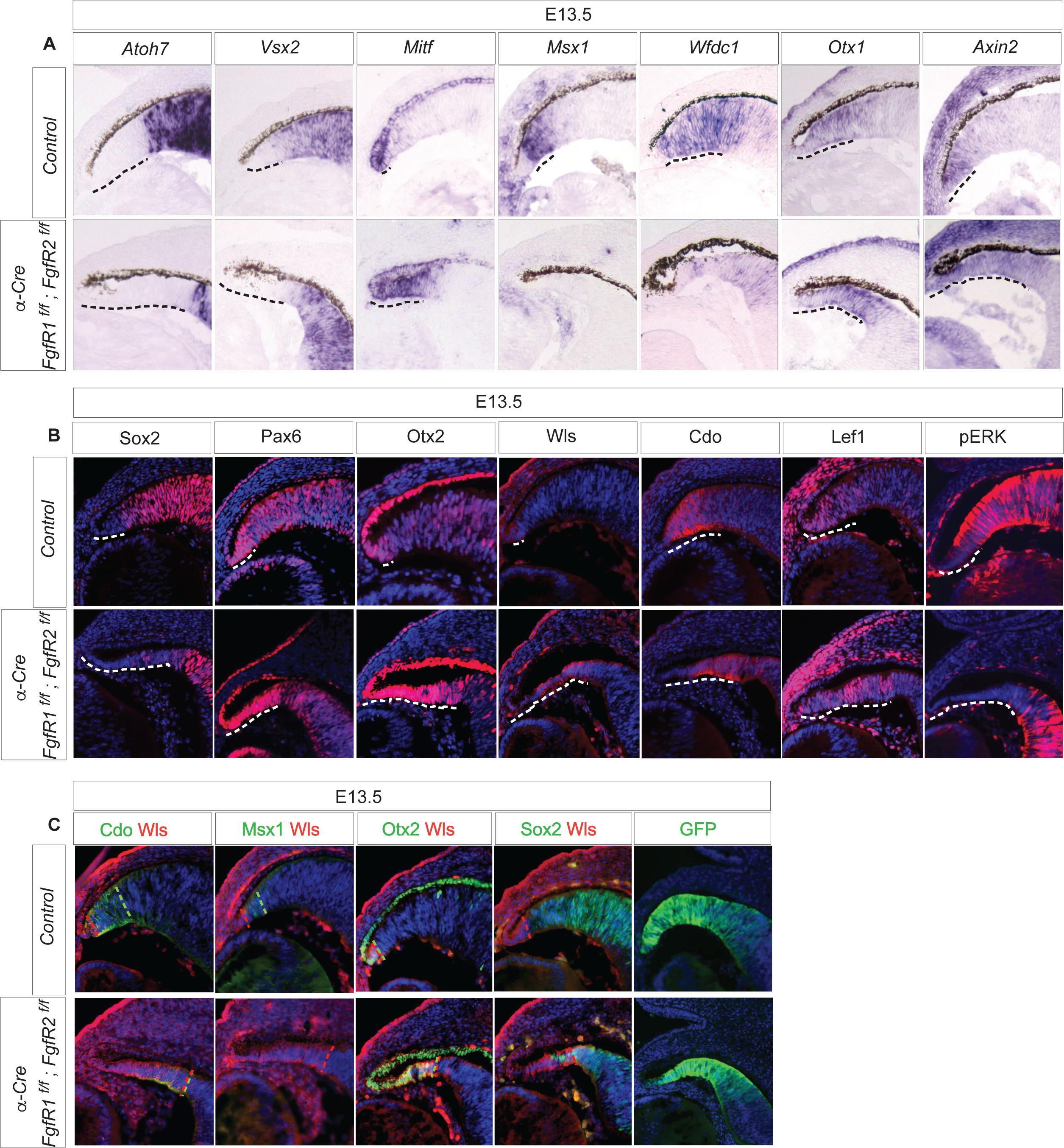
FGF signaling mutants lead to expansion of distal ciliary margin. **A.** Atoh7, a neural retinal marker indicates expansion of the ciliary margin in FGF signaling mutants. Vsx2 expression domain in the neural retina and proximal ciliary margin is also similarly ectopically pushed back into the neural retinal domain. Mitf expression in the RPE and distal ciliary margin is expanded ectopically towards the neural retinal domain. Msx1 expression in the proximal ciliary margin is noticeably lost. Wfdc1 expression from the proximal and medial ciliary margin is significantly diminished. Otx1 expression in the ciliary margin is expanded further towards the neural retina. Axin2 expression restricted normally to the distal and medial ciliary margin and a marker for Wnt signaling is expanded ectopically as well. **B.** Sox2, a neural retinal marker displays a graded expression in the proximal ciliary margin and is not present in the distal ciliary margin. It is noticeably absent in the proximal ciliary margin in the FGF signaling mutants. Pax6 expression is strong in the ciliary margin and weans towards the neural retina. The strong expression of Pax6 is seen expanded in FGF signaling mutants. Otx2 marks the RPE and distal ciliary margin cells. Otx2 expression is expanded into the neural retinal space. Wls expression in the RPE (which is not visible with pigmentation) and distal ciliary margin is also expanded. Cdo, a pan marker for ciliary margin is expanded towards the neural retinal space. Expression of Lef1 restricted normally to the distal and medial ciliary margin and a marker for Wnt signaling is expanded in the ectopic ciliary margin. pERK expression in the neural retina, proximal and medial ciliary margin is noticeably absent in the ciliary margin of FGF mutants. **C.** A comparison of Cdo, Msx1, Otx2, and Sox2 expression with Wls expression reveals the expansion of the distal ciliary margin at the expense of the proximal ciliary margin in the FGF mutants.

Atoh7 (Math5) is expressed exclusively in differentiating neural retinal population. In FGF mutants, the ciliary margin seems to extend ectopically towards the neural retina, thereby pushing back the Atoh7 expressing neural retinal population. Vsx2 (Chx10) is expressed in neural retinal progenitor cells and in the medial and proximal ciliary margin, but not the distal ciliary margin. In FGF mutants, the Vsx2 negative population of ciliary marginal cells expands, indicating the ectopic expansion of the distal ciliary margin. The expression of Otx1, a pan-marker of ciliary margin cells is also ectopically expanded towards the neural retinal space in FGF mutants. The expression of Msx1, a marker specific for the proximal ciliary margin is lost while the expression of Mitf, a gene expressed in the RPE and the distal tip of the ciliary margin is expanded in FGF mutants. Wfdc1 is expressed in the medial and proximal ciliary margin with a weak expression in neural retinal cells immediately adjoining the ciliary margin. Upon loss of FGF signaling, expression level of Wfdc1 is reduced. The expression of Axin2, a downstream effector of WNT signaling, which is strong in the distal ciliary margin and wanes towards the proximal ciliary margin is ectopically expanded in FGF mutants as well (Figure 2A). These gene patterns suggest that the loss of FGF signaling in the peripheral retina leads to an ectopic expansion of the peripheral tip of the RPE and ciliary margin, and a disruption in the compartmentalization of the ciliary margin cells in favor of distal ciliary margin cells. Using additional markers such as Sox2 (neural retinal and proximal ciliary margin marker), Pax6 (stronger expression in the ciliary margin), Otx2 (strong expression in the RPE and distal tip of the ciliary margin), Wntless (a G-protein coupled receptor that is essential for the secretion of Wnt ligands and is expressed in the RPE and the medial and distal tip of the ciliary margin) and Cdo (pan ciliary margin maker), we were able to confirm the expansion of the distal ciliary margin upon loss of FGF signaling in the peripheral retina. The expression of pERK – a readout of the FGF signaling cascade confirms the loss of FGF signaling in the ectopic ciliary margin; the expression of Lef1 – a transcriptional activator of WNT signaling shows a complimentary upregulation in the expanded ciliary margin (Figure 2B). To further qualitatively assess the contribution of the distal medial and proximal ciliary margin compartments in the FGF mutant, we analyzed the co-expression of cells expressing Wntless, with respect to other markers of ciliary margin (pan, medial, distal, neural retina+proximal) (Figure 2C). These results clearly demonstrate that the loss of FGF signaling in the peripheral retina leads to an expansion of the distal ciliary margin and a loss of the proximal ciliary margin.

Loss of FGF signaling in the peripheral retina leads to a reduction in the number of Ki67 and BrdU positive proliferative cells in the ectopic ciliary margin. The reduction in proliferation is most apparent at E13.5 (Figure S1). Previous studies have linked Hippo signaling to cell proliferation and differentiation at the ciliary margin. Indeed, we observed an expansion of YAP/TAZ expression in the mutant ciliary margin (Figure S1), suggesting increase in YAP/TAZ signaling as a result of loss of FGF signaling.

### The ciliary margin is induced by graded FGF signaling

To identify the ligands responsible for the activation and maintenance of FGF signaling at the neurogenic boundary, we conditionally deleted FGF9 and FGF3 using α-Cre. The loss of FGF9 resulted in a minor phenotype as seen by the expansion of the Mitf-expressing distal ciliary margin and Otx1 expression, similar to the loss of FGFR1 and FGFR2 (Figure 3A). The loss of both FGF9 and FGF3 aggravated the phenotype as seen by the additional reduction in Msx1 expression in medial ciliary margin cells. This shows that FGF9 and FGF3 are two of the possibly several important ligands in activating FGF signaling at the ciliary margin (Figure 3B). We also observe that loss of Mek1 and Mek2 phenocopy the loss of FGFR1 and FGFR2 (Figure 3C), consistent with MAPK pathway as the downstream mediator of FGF signaling in the ciliary margin.

**Figure 3:**
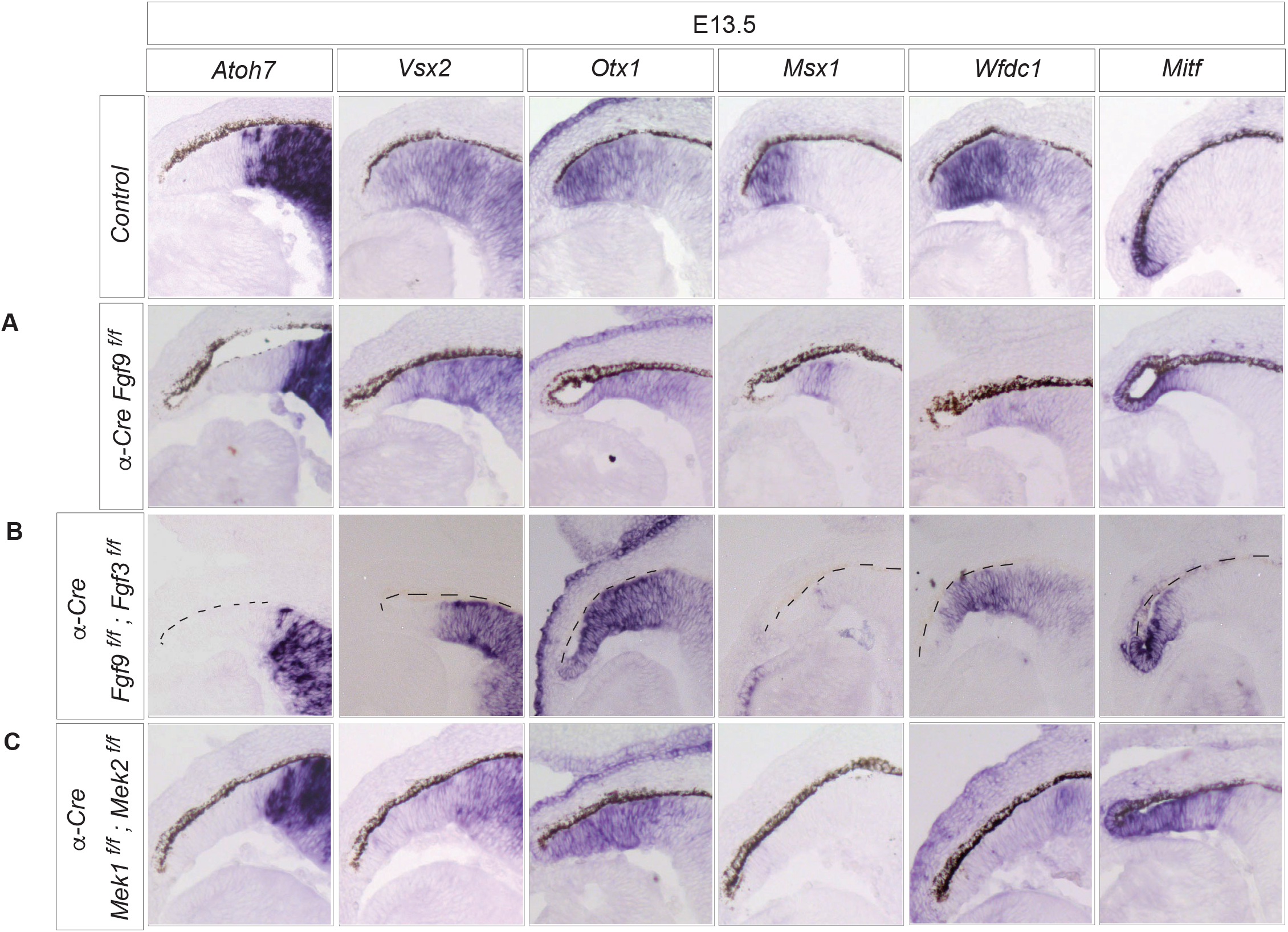
Fgf3 and Fgf9 signal via MAPK-ERK pathway to specify the ciliary margin. **A.** Loss of Fgf9 results in a slight expansion of Mitf expression and an equivalent expansion of the Otx1 expression domain. The expression of Atoh7 is also slightly expanded towards the neural retina. The expression of Vsx2, Msx1 and Wfdc1 remains unaltered. **B.** Loss of both Fgf3 and Fgf9 results in a similar phenotype as the loss of Fgf9 with additional reduction in Msx1 expression in the ciliary margin. **C.** Loss of Mek1 and Mek2 phenocopies the loss of FGFR1 and FGFR2.

According to our model and results from Figure 2, FGF signaling is required for development of the proximal and medial ciliary margin compartments. To demonstrate the sufficiency of FGF signaling for the proximal and medial ciliary margin, we first constitutively expressed Fgf8 using α-Cre (Figure 4A). As expected, over-expression of Fgf8 results in a near complete trans-differentiation of the RPE into the neural retina. The trans-differentiated RPE expresses Atoh7 and Vsx2 but not Mitf, indicating its conversion to the neural retina. Next, we reduced the dosage of Fgf8 signaling by deleting Fgf receptors 1 and 2, leaving only Fgf receptors 3 and 4 intact (Figure 4B). As indicated by the expression of FGF downstream response gene Spry2, Fgf8 overexpression alone led to greatly elevated FGF signaling in the transformed RPE, which was significantly attenuated after the removal of Fgfr1 and 2. Strikingly, these Fgfr1/2-deficient cells do not express Atoh7, Vsx2 or Wfdc1, but express Mitf, Msx1 and Otx1, indicating a mixture of distal and proximal ciliary margin cells. This is accompanied by increased expression of Axin2, a downstream response gene to Wnt signaling. Therefore, a low level of Fgf signaling induces the ciliary margin fate with moderate Wnt signaling activity.

**Figure 4:**
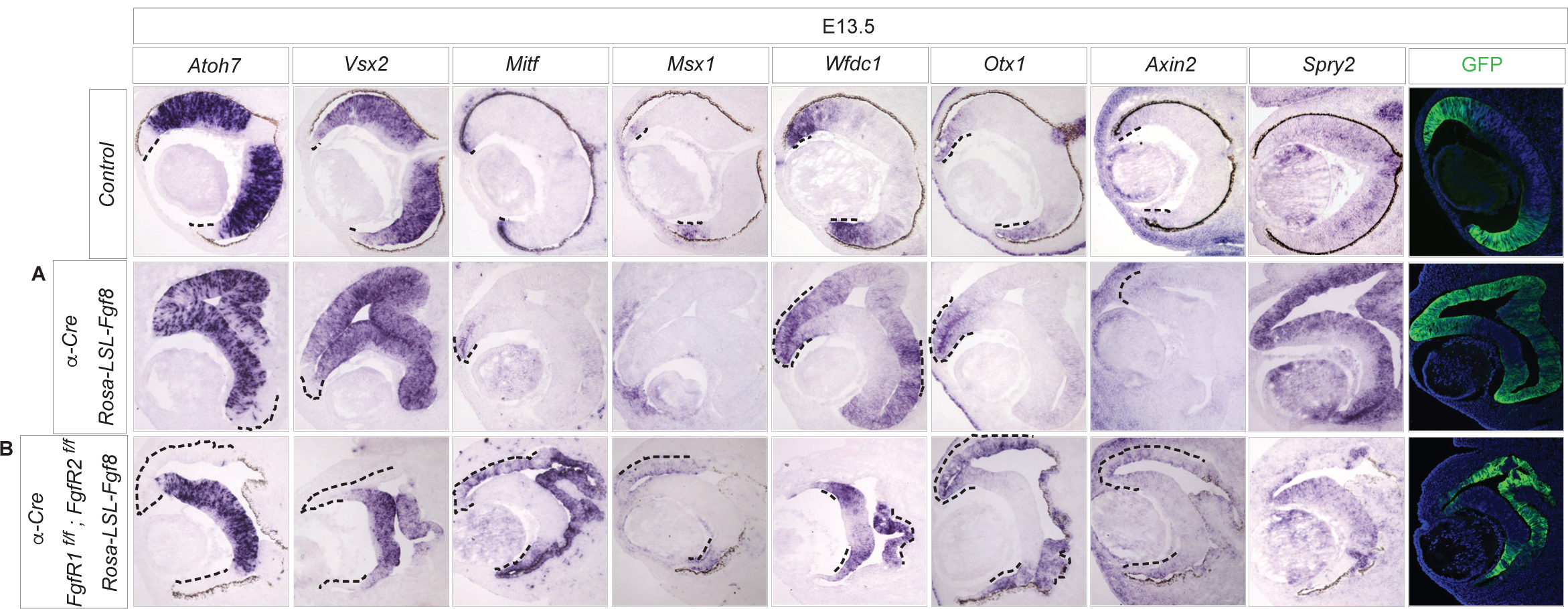
A titration of FGF signaling is needed for the specification of the different ciliary margin zones. **A.** Constitutive expression of Fgf8 driven by α-Cre leads to the trans-differentiation of the RPE into neural retina. Atoh7 and Vsx2 is expressed in the double-layered neural retina. Trans-differentiated RPE does not express Mitf and Msx1. Wfdc1 is strongly expressed in the optic cup rim where Otx1 is also expressed indicating the residual ciliary margin. Axin2 is expressed weakly in the optic cup rim and is not seen elsewhere in the trans-differentiated retina indicating that a high level of Fgf signaling is incompatible with Wnt signaling. **B.** A titrated effect of Fgf8 elicited by the removal of FGFR1 and FGFR2 while constitutively expressing Fgf8 produces a partial trans-differentiation of the RPE into ciliary margin. Atoh7 and Vsx2 expression is restricted to the inner neuro-epithelium. The partially trans-differentiated RPE consisting of non-pigmented cells express Mitf, Msx1 and Otx1 but not Wfdc1.

### Wnt ligands from the surface ectoderm activate WNT signaling to specify the ciliary margin

Previous studies have shown that loss of β-catenin, which has bipartite functions in both canonical WNT signaling and cell adhesion, disrupted ciliary margin development (Liu et al., 2007). Here, we further characterize dysgenesis of the ciliary margin using the markers highlighted above. Conditional deletion of β-catenin in the peripheral retina using α-Cre results in a disrupted neurogenic boundary in certain aspects complimentary to that of the loss of FGF signaling (Figure 5A). The expression of Atoh7 and Vsx2 is expanded into the presumptive ciliary marginal space, indicating the ectopic expansion of the neural retinal domain. The expression of Otx1 is largely restricted to a few cells at the very distal tip of the optic cup. The expression of Msx1 in proximal ciliary margin cells is absent. The expression of Wfdc1 is also restricted to a few cell layers at the distal tip of the optic cup and the expression of Mitf and Axin2 is restricted only to the RPE. These results indicate that a minimal ciliary margin at the distal tip of the optic cup, likely a result of incomplete gene deletion. Using a second set of redundant markers, we confirm that indeed the neural retina is expanded into the presumptive ciliary margin space (Sox2 and Pax6 expression), leaving only a few ciliary margin cells intact (Otx2, Wls and Cdo expression). Importantly, loss of canonical WNT signaling (Lef1 expression) in the peripheral retina is accompanied by expansion of FGF signaling (pERK expression) in the presumptive ciliary margin (Figure 5B). Since the loss of β-catenin also affects the cell-cell adhesion properties, we conditionally deleted Lrp5 and Lrp6 – two key receptors specific to the canonical WNT pathway (Figure 5A). We show that knockout of Lrp5 and Lrp6 phenocopy β-catenin deletion in the peripheral retina, which translates to loss of ciliary body architecture in the adult eye (Fig 5 A and B). These results confirm the essential requirement of canonical WNT signaling in specifying the ciliary margin.

**Figure 5:**
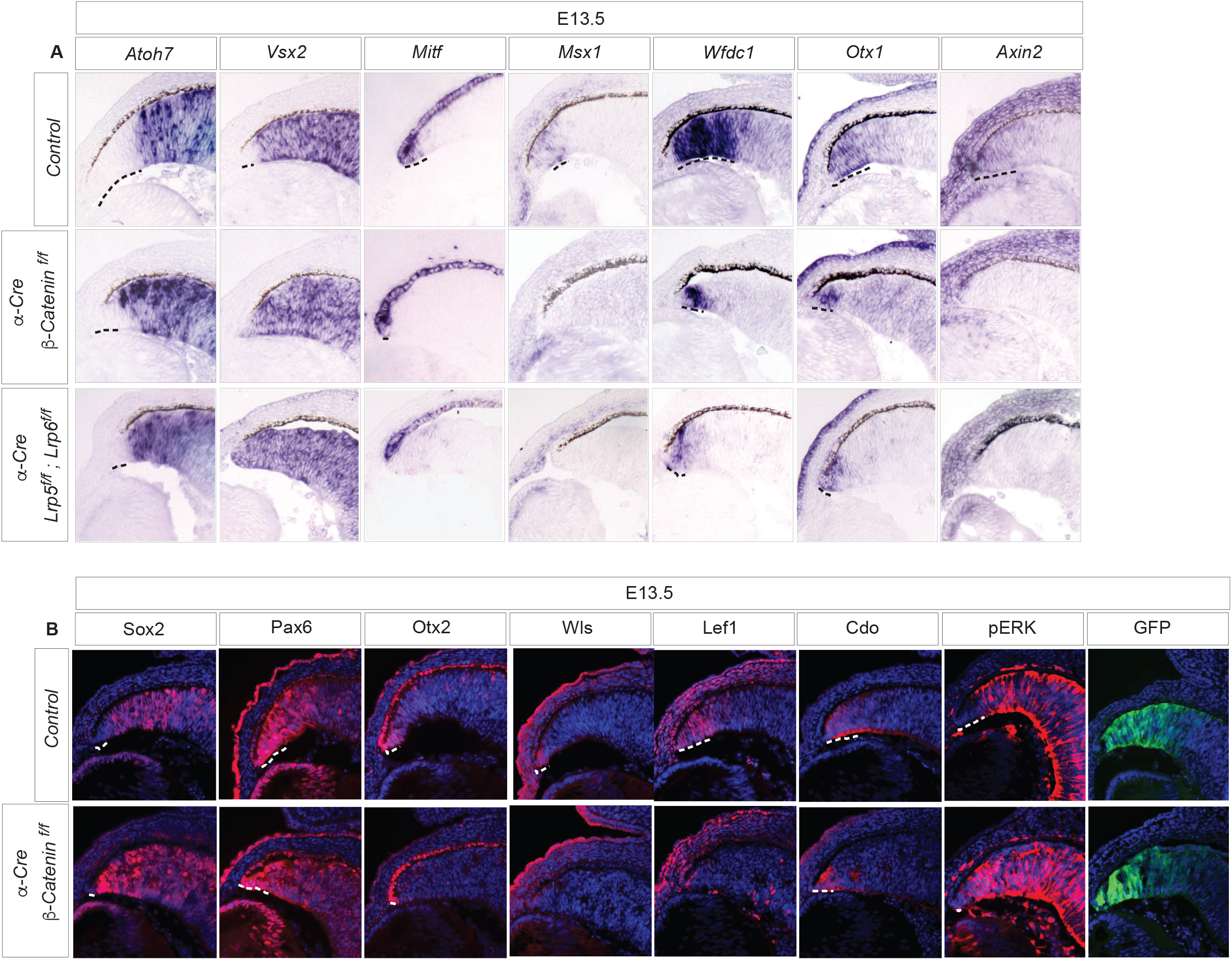
Canonical WNT signaling mutants leads to loss of distal and medial ciliary margin. **A.** Loss of either β-catenin or Lrp5/Lrp6 in the peripheral retina resulted in similar defect in the ciliary margin. Atoh7 expression domain is pushed closer towards the distal tip of the optic cup indicating ectopic expansion of the neural retinal domain. Vsx2 expression is expanded to the distal tip of the optic cup. Mitf expression in the distal ciliary margin and Msx2 expression in the ciliary margin are lost in WNT signaling mutants. Wfdc1 and Otx1 expression is restricted to a small portion of the distal ciliary margin while Axin2 expression is completely lost from the ciliary margin. **B.** Stronger expression of Sox2 is seen in the distal tip of the ciliary margin in the WNT signaling mutants. The stronger expression of Pax6 in the ciliary margin is restricted to the distal tip of the optic cup. Otx2 expression from the distal ciliary margin is lost, Wls expression from the distal ciliary margin is lost and Lef1 expression from the distal and medial ciliary margin is lost. Cdo expression is restricted to a small region of the distal optic cup. pERK expression is expanded towards the distal tip of the ciliary margin.

We next determine the source of WNT ligands that elicit a WNT response in the ciliary margin. During development, the expression of Wnt ligands is typically seen in the developing lens, RPE, and periocular mesenchyme but not the retina (Carpenter et al., 2015; Fuhrmann, 2008). We hence reasoned that WNT signaling is non-cell autonomously activated by paracrine Wnt ligands produced elsewhere. Previous studies have shown that loss of Wntless from the developing surface ectoderm leads to a trans-differentiation of the peripheral RPE into neural retina (Carpenter et al., 2015). We asked if the loss of Wntless, and hence the secretion of Wnt ligands from the surface ectoderm also leads to the loss of ciliary margin markers (Figure 6). Indeed, the hinge region at the junction of the neural retina and the RPE does not express the ciliary margin markers Mitf and Msx1, although a few cells do express Otx1 and Wfdc1, indicting the minimal presence of proximal ciliary margin. This phenotype is reminiscent of the loss of β-catenin and hence that of canonical WNT signaling in the peripheral retina. These results show that WNT signaling at the peripheral retina requires Wnt ligands originating from the developing lens/surface ectoderm. To determine if the periocular mesenchyme acts as a relay between the surface ectoderm and the ciliary margin, we next deleted Wntless from the periocular mesenchyme using Wnt1-Cre. However, we were unable to identify any morphological or phenotypical changes in the ciliary margin upon loss of Wnt ligands from the developing periocular mesenchyme. Taken together, these results demonstrate the surface ectoderm-derived Wnt ligands are required for the ciliary margin formation.

**Figure 6:**
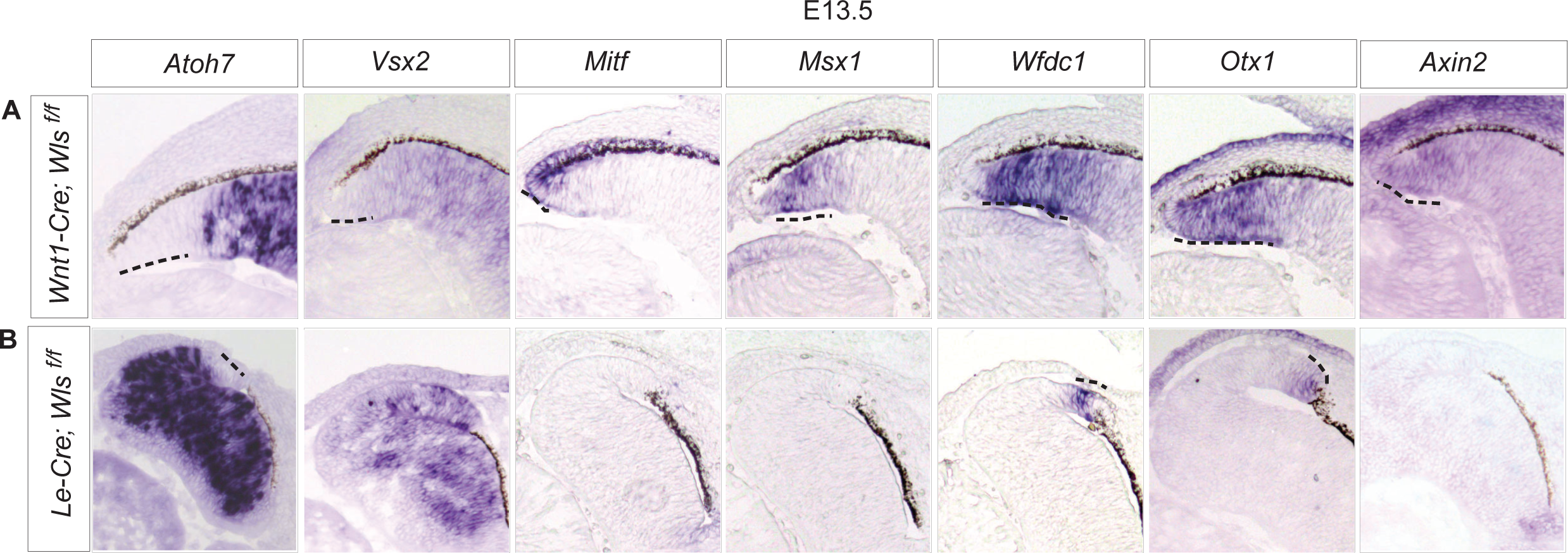
Wnt ligands from the developing lens/surface ectoderm activate Wnt signaling at the ciliary margin. **A.** Deletion of Wntless from the periocular mesenchyme using Wnt1-cre does not cause any phenotypic changes in neural retinal and ciliary margin marker expression. **B.** Deletion of Wntless specifically from the developing surface ectoderm using Le-Cre leads to trans-differentiation of the peripheral RPE into neural retinal like cells. Atoh7 and Vsx2 expression is seen in the trans-differentiated cells while expression of Mitf and Msx1 is lost. Wfdc1 and Otx1 are expressed in a small group of cells indicating the presence of proximal ciliary marginal cells. Axin2 expression is completely lost from the developing lens, surface ectoderm and ciliary margin.

### Combined activation of both FGF and WNT signaling induces the trans-differentiation of the RPE into ciliary margin

To examine the interaction between FGF and WNT signaling, we first constitutively expressed only Wnt1 gene driven by α-Cre (Figure 7). Interestingly, we did not see any changes in the ciliary margin or neural retina. We then over-expressed both Fgf8 and Wnt1 genes. We see a similar expansion of the RPE into a multi-layer structure as with the constitutive expression of only Fgf8. As expected, these cells displayed elevated FGF signaling as indicated by strong expression of Spry2, but they also maintain active Wnt signaling as demonstrated by the robust expression of Lef1. Intriguingly, they also express Vsx2 but not Math5, suggesting that they are not transformed into neural retina. Instead, there are prevalent expression of Mitf and Otx1 and restricted expression of Msx1. These results indicate that the RPE can be transformed into ciliary margin by simultaneous activation of both FGF and WNT signaling.

**Figure 7:**
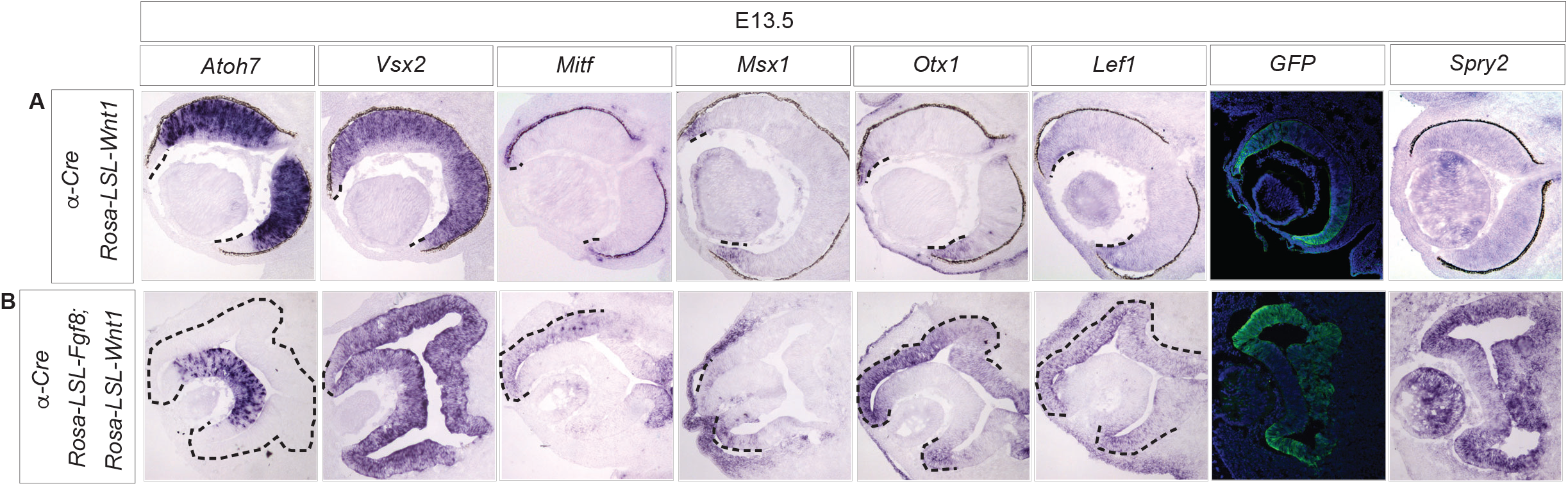
Constitutive activation of both Fgf and Wnt signaling induces trans-differentiation of the RPE into ciliary margin. **A.** Constitutive expression of Wnt1 driven by α-Cre does not elicit any phenotypic changes in the expression of neural retinal or ciliary margin marker expression. **B.** Constitutive expression of both Fgf8 and Wnt1 driven by α-Cre induces trans-differentiation of the RPE, which however do not express Atoh7. The cells express Chx10, Mitf and Otx1, while also faintly expressing Msx1, indicating the presence of a mixture of both proximal and distal ciliary margin cells. Lef1 expression is also seen in the trans-differentiated layer indicating activation of Wnt signaling.

## DISCUSSION

The ciliary margin is a crucial developmental structure for development of the ciliary body and iris. The ciliary margin at the peripheral retina is not morphologically distinct from the adjacent neural retina. However, molecular differences between these two cell populations lead them to adopt different cell fate. Previous studies have illuminated the essential role of WNT signaling in development of the ciliary margin, and how the boundary between the neurogenic retina and ciliary margin is maintained (Heavner et al., 2014; Liu et al., 2007). Although FGF signaling has been implicated in development of the ciliary margin in chick (Dias da Silva et al., 2007), the mechanistic intersection of FGF and WNT signaling at the peripheral retina has not been previously studied. In this study, we have successfully shown that the compartmentalization of the ciliary margin into proximal (closer to FGF-high neural retina), medial, and distal (closer to WNT-high RPE) is specified by the intersection of FGF and WNT gradients. Using mouse genetics, we show that FGF signaling in the proximal part of the ciliary margin inhibits WNT signaling while WNT signaling in the distal part of the ciliary margin inhibits FGF signaling and that both FGF and WNT signaling systems are needed to specify the ciliary margin. We also show that a low level of FGF signaling is needed for the specification of the ciliary margin while a high level of FGF signaling leads to the development of the neural retina. Very importantly, these results explain how the boundary between the neural retina and the ciliary margin is created by a careful titration of FGF and WNT signaling.

This study demonstrated the requirement of FGF signaling in development of the entire ciliary margin. We showed that in the absence of FGF signaling, the distal ciliary margin is expanded ectopically into the neural retinal domain. Using constitutively active Fgf8, we have also demonstrated the need for a gradient of FGF signaling in specifying the different compartments of the ciliary margin. Loss of FGF signaling in the peripheral retina also leads to aniridia in the adult eye resulting from the disruption in ciliary margin development. We have determined the upstream and downstream components of the FGF signaling cascade in development of the ciliary margin. Several studies have shown that Fgf ligands are expressed in the retina, lens and periocular mesenchyme during retinal development (de Iongh and McAvoy, 1993; Kurose et al., 2005; McMahon et al., 2009; Vogel-Hopker et al., 2000; Zhao et al., 2001). In this study, we were able to demonstrate that pERK expression is high in the neural retina and diminishes towards the peripheral optic cup in a gradient-like manner. Conditional deletion of Fgf9 using α-Cre had a small but significant effect on the compartmentalization of the ciliary margin. Specifically, the distal ciliary margin was slightly expanded into the neural retinal domain. A similar effect was seen in a previous study of Fgf9 systemic knockout (Zhao et al., 2001). Our study also showed an additive effect of deleting Fgf3 along with Fgf9 on the compartmentalization of the ciliary margin. We also place Mek1 and Mek2 in the FGF signaling pathway necessary for the development of the ciliary margin. These results demonstrate FGF-MAPK as the signaling cascade important for development and compartmentalization of the ciliary margin.

Previously, β-catenin had been demonstrated to have a critical role in development of the ciliary margin (Liu et al., 2007). Since β-catenin has dual roles in cell-cell adhesion and as a transcriptional activator of canonical WNT signaling (Brembeck et al., 2006), we conditionally both Lrp5 and Lrp6 to demonstrate that the canonical WNT signaling indeed plays an important role in development of the ciliary margin. We also showed that loss of WNT signaling leads to expansion of FGF signaling in the distal optic cup. These results together demonstrated the antagonistic interactions of FGF and WNT signaling systems. Using a combined constitutive activation of Fgf8 and Wnt1, we showed that indeed both signaling systems needed to intersect to specify the medical ciliary margin. This provides for a dose-based gradient model to explain development of the ciliary margin. We also identify the developing lens/surface ectoderm as the key source of WNT ligands for development of the ciliary margin (Carpenter et al., 2015). In our study, we showed that while the neural retina expands into the presumptive RPE space, the ciliary margin is not specified at the junction of the neural retina and RPE in the absence of secreted Wnt ligands from the lens. In concurrence with a previous study (Carpenter et al., 2015), we also found that preventing secretion of Wnt ligands from the periocular mesenchyme does not affect development of the ciliary margin. Our studies also looked at the loss of Wntless from the peripheral retina (data not shown) and found no phenotypic differences in the development of ciliary margin. These results show that the Wnt ligands from the surface ectoderm likely signal to the optic cup directly without a relay intermediary.

## MATERIALS AND METHODS

### Animals

Mice carrying *R26^Fgf8^,Mek1^flox^* and *Mek2^-/-^* alleles were bred and genotyped as described (Lin et al., 2013; Newbern et al., 2008). *Fgfr2^flox^* line was obtained from Dr. Dr. David Ornitz (Washington University Medical School), Pax6 α-Cre (α-Cre) from Dr. Ruth Ashery-Padan (Tel Aviv University), *Fgf3 ^flox^* from Dr. Suzanne L Mansour (University of Utah), *R26-Fgf8^LSL^* from Dr. Yiping Chen (Tulane University), *R26-Wnt1^LSL^* from Dr. Thomas Carroll (UT Southwestern), and *Fgfr1^flox^* and *Wntless ^flox^* from Jackson lab (Ashery-Padan et al., 2000; Belanger et al., 2003; Bissonauth et al., 2006; Brault et al., 2001; Carpenter et al., 2010; Carroll et al., 2005; Danielian et al., 1998; Hoch and Soriano, 2006; Joeng et al., 2011; Lin et al., 2013; Lin et al., 2006; Marquardt et al., 2001; Su et al., 2010; Tuveson et al., 2004; Urness et al., 2010; Weinstein et al., 1998; Yu et al., 2003).

Mice were maintained on a mixed genetic background. At least three animals were analyzed for each of the crosses described (except for the Rosa-LSL-Fgf8-Wnt1 only 2 animals were analyzed). We did not observe phenotypic variations between Cre-heterozygous controls and no-Cre homozygous controls. These two genotypes are hence together described as controls. All animal procedures were performed according to the protocols approved by the Columbia University’s Institutional Animal Care and Use Committee.

### Histology

Mice were time-mated and the noon of the day vaginal plug was first visualized was designated E0.5. Embryos were harvested in ice-cold PBS, fixed in 4% PFA overnight, equilibrated in 30% sucrose and cryo-embedded in OCT. Samples were sectioned in a cryostat at 10 μm thickness and processed for further staining. Sections were stained with Hematoxylin and Eosin (H&E) for gross morphological analysis.

### Immunohistochemistry

Cryo-sections were processed for immunohistochemistry using previously established protocols. Briefly, slides were washed in PBS 3 times for 5 minutes each. This was followed by PBST (0.3% Triton) washes 3 times for 5 minutes each. Sections were then blocked in 10% Horse Serum in PBST (blocking buffer) for 1 hour. Primary antibodies listed in Table 2 were diluted in blocking buffer according to the dilutions and added to the slides and placed in 4C overnight. If antigen retrieval was performed, it was done so using 10mM Sodium Citrate 0.05% Tween, pH 6.0 (Antigen retrieval buffer). Briefly, slides were washed in PBS 3 times for 5 minutes each. Slides in antigen retrieval buffer were placed in an antigen retriever (Aptum Biologics) for 3 cycles of heating. Slides were then cooled to room temperature for an hour before continuing with the PBST washes, blocking and addition of primary antibody as described above. Alexa Fluor secondary antibodies was added the next day for 2 hours at room temperature followed by a short incubation of 5 minutes with DAPI. Slides were cover-slipped using NPG-Glycerol mounting medium for imaging. Antibodies used were: Goat anti-Sox2 (Santa Cruz Biotechnology), Rabbit anti-Pax6 (Covance Research Products Inc.), Goat anti-Otx2 (Santa Cruz Biotechnology), Rabbit anti-Wls (Seven Hills Bioreagents), Goat anti-Cdo (R&D Systems), Rabbit anti-Lef1 (Cell Signaling Technology), Rabbit anti-pERK (Cell Signaling Technology), Goat anti-Msx1 (R&D Systems), Rabbit anti-GFP (Thermo Fisher Scientific), Chick anti-GFP (Aves Labs), Mouse anti-Brdu (Developmental Studies Hybridoma Bank) and Mouse anti-αSMA (Sigma-Aldrich).

### In-situ hybridization

Cryo-sections were processed for in-situ hybridization using previously established protocols. Briefly, DIG-labeled mRNA *in situ* probes diluted in hybridization buffer was added to the slides and incubated at 65C overnight. High stringent buffers and MABT washes were done before blocking the slides with 10% Goat Serum in MABT. Anti-DIG antibody diluted in MABT (with 1% Goat Serum) was added to the slides and incubated at 4C overnight. Slides were then washed in MABT followed by Alkaline Phosphatase buffer (NTMT). BM Purple (Millipore Sigma) was added to the slides and the color reaction was allowed to develop overnight. Slides were then briefly washed in PBS and cover-slipped with mounting medium for imaging. Probes used were obtained from: Atoh7 from Dr. Tom Glaser (University of California Davis), Vsx2 from Dr. Roderick McInnes (McGill University), Mitf from Dr. Hans Arnheiter (NIH), Msx1 from Dr. Valerie Wallace (University of Toronto), Wfdc1 from Dr. Jean Hebert (Albert Einstein College of Medicine), Otx1 from Dr. Valerie Wallace (University of Toronto), Lef1 from Dr. Valerie Wallace (University of Toronto), and Axin2 from Dr. Frank Costantini (Columbia University).

### BrdU labeling

Stock concentrations of BrdU was prepared at 10mg/ml and stored in −20C. Animals were weighed and injected with 100mg/kg BrdU intraperitoneally. Animals were euthanized 2-3 hours after injection and embryos/tissue harvested in ice-cold PBS and fixed in 4%PFA briefly for 1 hour. 30% Sucrose equilibrated samples were cryo-embedded and sectioned. Antigen retrieval for BrdU was done by incubating slides in 2N HCl for 20 minutes prior to blocking.

## ACKNOWLEDGEMENTS

The authors thank Drs. David Ornitz, Ruth Ashery-Padan, Suzzane L Mansour, Yiping Chen, Thomas Carroll and Richard Lang for mice and Drs. Tom Glaser, Roderick McInnes, Hans Arnheiter, Valerie Wallace, Jean Hebert and Frank Costantini for *in situ* hybridization probes. We thank John Peregrin for technical support, and members of the Zhang lab for helpful discussions and comments. We also thank Dr. Carol Mason for critical reading of the manuscript.

This work was supported by the National Institutes of Health (EY017061, EY018868 and EY 025933 to XZ). The Columbia Ophthalmology Core Facility are supported by NIH Core grant 5P30EY019007 and unrestricted funds from Research to Prevent Blindness (RPB). RB is a recipient of the Knights Templar Eye Foundation Career-Starter Grant. CT is a recipient of Jonas Scholar award. XZ is supported by Jules and Doris Stein Research to Prevent Blindness Professorship.

---

**Supplemental Figure 1:**
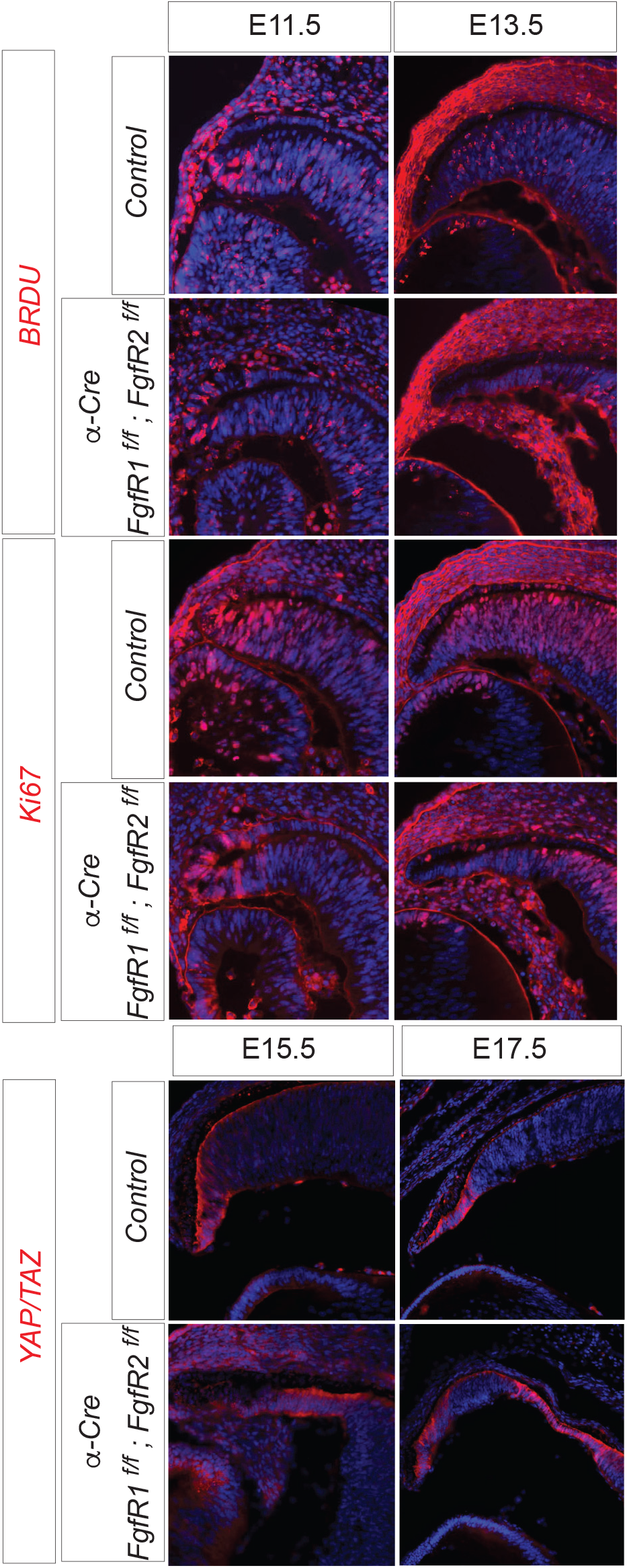
Loss of FGF signaling in the peripheral retina affects proliferation of developing ciliary margin cells. A. Brdu and Ki67 labeling was reduced in the ciliary margin upon loss of FGF signaling at E11.5 and E13.5. B. Strong YAP/TAZ expression at E15.5 and E17.5 is seen in the ciliary margin. The expression domain of YAP/TAZ is expanded.

## Notes

The authors declare that there is no conflict of interest with regard to the manuscript.

